# Alterations in genes associated with cytosolic RNA sensing in whole blood are associated with coronary microvascular disease in SLE

**DOI:** 10.1101/2024.02.19.580713

**Authors:** Lihong Huo, Erica Montano, Gantseg Tumurkhuu, Moumita Bose, Daniel S. Berman, Daniel Wallace, Janet Wei, Mariko Ishimori, C. Noel Bairey Merz, Caroline Jefferies

**Author notes:** Corresponding Author: Caroline Jefferies, PhD, Kao Autoimmunity Institute, 121 N San Vincente Blvd., Los Angeles, California, 90211. The authors have no disclosures to report.

## Abstract

**Objective:** To investigate whether gene signatures discriminate systemic lupus erythematosus (SLE) patients with coronary microvascular dysfunction (CMD) from those without and whether any signaling pathway is linked to the underlying pathobiology of SLE CMD.

**Methods:** This study collected whole blood RNA samples from female subjects aged 37 to 57, comprising 11 SLE patients (4 SLE-CMD, 7 SLE-non-CMD) and 10 HC. Total RNA was then used for library preparation and sequencing. Differential gene expression analysis was performed to identify gene signatures associated with CMD in SLE patients using DEseq2 v1.42.0. Gene Set Enrichment Analysis were performed by ClusterProfiler v4.10.0 and pathfindR v2.3.1.

**Results:** RNA-seq analysis revealed 143 differentially expressed (DE) genes between the SLE and HC groups. GO analysis indicated associations with virus defense and interferon signaling in SLE. 14 DE genes were identified from comparison between SLE-CMD and SLE-non-CMD with adjusted parameters (padj < 0.1). Notably, SLE-CMD exhibited elevated levels of genes associated with RNA sensing, while downregulated genes in SLE-non-CMD were associated with blood coagulation and cell-cell junction. Further investigation highlighted differences in IFN signaling and ADP-ribosylation pathways between SLE-CMD and SLE-non-CMD, suggesting distinct molecular mechanisms underlying vascular changes in CMD and reduced left ventricular function in non-CMD.

**Conclusion:** Our study identified a unique gene signature in SLE-CMD compared to the HC group, highlighting the significant involvement of type 1 interferon, RIG-I family proteins, and chronic inflammation in the progression of SLE-CMD. The intricate relationship between SLE-CMD and these factors underscores their probable role in initiating and advancing SLE-CMD.

## Introduction

Patients with SLE were over three times more likely to die of cardiovascular disease than patients in the general population[1]. Many women with SLE frequently report chest pain in the absence of obstructive coronary artery disease (CAD) due to coronary microvascular dysfunction (CMD), a form of ischemia with no obstructive CAD. CMD is a heart condition where blood flow response is impaired, resulting in reduced coronary flow reserve (CFR)[2]. This is associated with increased resistance in small blood vessels, spasms, limited myocardial perfusion reserve, and potential heart muscle ischemia, despite minimal blockage in the main heart arteries (less than 50% narrowing or a fractional flow reserve over 0.80)[3]. In terms of risk factors for CMD in the general population, notable associations have been found in women who have impaired coronary flow reserve with age, hypertension, smoking history, elevated heart rate, and low HDL[4]. Chronic inflammation also plays an important role in the pathophysiology of CMD[5]. For example, elevated levels of CRP, VCAM-1, PAI-1, vWF have been reported in CMD patients[6]. However, the molecular mechanisms underlying the development of CMD in SLE patients is unknown.

Cardiac magnetic resonance imaging (cMRI) is a non-invasive technology that allows for analysis of cardiac function, including CMD. cMRI measures determines the mass and volumes of the heart, in addition to providing structural imaging of the myocardial tissue to detect fibrosis. Stress perfusion imaging allows measurement of coronary blood flow, which permits detection of ischemia and valve disorders. The semi-quantitative myocardial perfusion index (MPRI) is both sensitive and specific for the diagnosis of CMD (a MPRI of less than 1.84 being diagnostic of CMD), whereas measures of left ventricular diastolic volumes and ejection fraction detect cardiac dysfunction. Previous studies by our group have demonstrated that approximately 40% of SLE patients have CMD on cMRI [7-9]. Interestingly in our most recent study we compared cardiac function by cMRI with clinical and inflammatory markers in a cohort of 13 SLE women[9]. We found that left ventricular function and cardiac strain were impaired in patients with SLE compared to reference controls and correlated with increased SLICC damage index and CRP levels. However, when we analyzed cardiac function and markers of inflammation in patients with and without CMD, we observed that patients without CMD contributed more to the observed differences between SLE and reference control groups, perhaps driven by increased left ventricular mass in patients without CMD versus patients with CMD. This prompted us to apply additional analyses to gain a better understanding of the molecular mechanisms underpinning CMD in SLE. RNA-sequencing (RNA-seq) offers valuable insights into the gene expression patterns within the transcriptome. In this study, our objective was to evaluate whether distinctive gene signatures can differentiate between SLE patients with CMD (SLE-CMD) and those without (SLE-non-CMD) by examining gene expression in healthy controls, SLE-CMD, and SLE-non-CMD whole blood. The findings presented in this article establish connections between peripheral biomarkers and the underlying pathobiology of SLE-CMD.

## Materials and Methods

### Patients and Healthy Controls

This cross-sectional study was approved by the institutional review board at Cedars-Sinai Medical Center, and all participants gave informed consent prior to participation. Participants were recruited from the lupus clinic. Inclusion criteria consisted of female SLE subjects (aged 37 to 57 years at baseline) with chest pain due to suspected angina. Exclusion criteria included documented obstructive CAD and contraindications to coronary computed tomography angiography (CCTA) or cardiac magnetic resonance imaging (cMRI). Healthy controls were age matched and recruited from Cedars-Sinai Medical Center. blood samples were collected from all patients and controls in the fasting state using PAXgene blood RNA tubes.

### PAXgene RNA isolation

Blood samples were kept in PAXgene RNA tubes in -80 until ready to process. RNA was isolated using the PAXgene Blood RNA kit according to the manufacturers guidelines (PreAnalytiX). Eluted RNA was dissolved in RNase-free water. The quality and quantity of RNA were evaluated using the Agilent 2100 BioAnalyzer (Santa Clara, CA, USA).

### RNA Sequencing

Total RNA samples were analyzed for RNA integrity on the 2100 Bioanalyzer using the Agilent RNA 6000 Nano Kit (Agilent Technologies, Santa Clara, CA) and quantified using the Qubit RNA HS Assay Kit (ThermoFisher Scientific, Waltham, MA). Three hundred ng of total RNA Total was ribodepleted using the RiboCop Depletion Kit Human/Mouse/Rat v2 (Lexogen Inc., Greenland, NH). Stranded RNA-Seq library construction was performed using the xGen Broad-Range RNA Library Prep Kit (Integrated DNA Technologies, Coralville, IA). Library concentration was measured with a Qubit fluorometer (ThermoFisher Scientific), and library size was evaluated on a 4200 TapeStation (Agilent Technologies). Multiplexed libraries were sequenced on a NovaSeq 6000 (Illumina, San Diego, CA) using 75bp single-end sequencing. On average, approximately 50 million reads were generated from each sample.

### Bioinformatics and data analysis

Raw reads obtained from RNA-Seq were aligned to the transcriptome using STAR (version 2.5.0)/RSEM (version 1.2.25) with default parameters, using a custom human GRCh38 transcriptome reference downloaded from https://www.gencodegenes.org, containing all protein coding and long non-coding RNA genes based on human GENCODE version 33 annotation. Expression counts for each gene in all samples were normalized by a modified trimmed mean of the M-values normalization method and the unsupervised principal component analysis (PCA) was performed with DESeq2 Bioconductor package version 1.42.0 in R version 4.3. Each gene was fitted into a negative binomial generalized linear model, and the Wald test was applied to assess the differential expressions between two sample groups by DESeq2. Benjamini and Hochberg procedure was applied to adjust for multiple hypothesis testing, and differential expression gene candidates were selected with a false discovery rate less than 0.05. For functional enrichment analysis across sample groups, we conducted genes enrichment analysis using the R package “clusterProfiler v4.10.0”[10] and “pathfindR v2.3.1[11].

### Results

Clinical information of the study subjects is described in our previous analysis[9]. All study subjects are female, with no significant differences in age or subject characteristics (such as BMI, fasting glucose or insulin levels, and inflammatory markers such as C3, C4, CRP and ESR) between groups. Whole blood transcriptomes from the cohort of 11 SLE patients (4 with CMD and 7 without CMD) and 10 age matched healthy controls (HC) were analyzed by RNA-sequencing (RNA-seq).

Principal Component Analysis (PCA) unveiled distinctive expression profiles between HC and SLE (Figure 1A). After data normalization using DEseq2, preprocessing and filtering with the criteria of padj <0.05, We identified 143 differentially expressed (DE) genes when comparing the SLE group and HC group. The DE gene analysis delineated a discernible molecular signature between SLE and HC, as displayed in Figure 1B. Upon further filtering with a |log2FC|>0.5 threshold, we identified 52 upregulated genes and 50 downregulated genes. Among the top 10 upregulated genes were IFI27, IFI44L, RSAD2, IFI44, SIGLEC1, IFIT1, SLC12A1, RPL23P3, CTXN2, ISG15, while the top 10 downregulated genes were RN7SKP227, RN7SL1, NTN4, RN7SL653P, SLC1A7, SGCD, S1PR5, KIR2DL3, LIM2, MMP23B. To gain insights into the functionality of those genes, we conducted a comprehensive Gene Ontology (GO) analysis. As expected, GO analysis results revealed SLE is significantly associated with gene enrichment of functions related to defense response to virus, type I interferon signaling pathway, and response to virus in upregulated DE genes (Figure 1C)[12-15].

**Figure 1.**
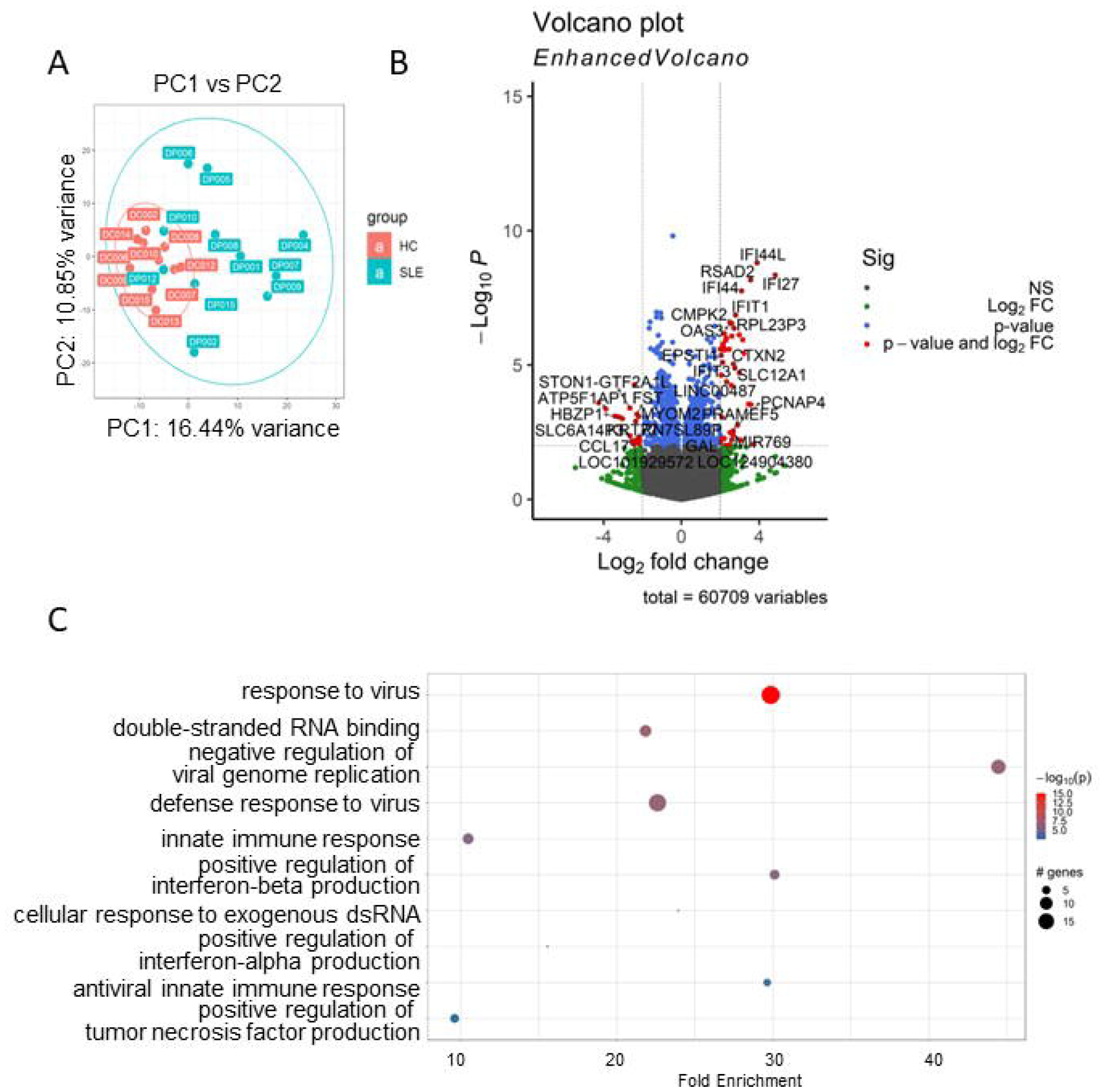
DEG and pathway analysis of SLE vs HC whole blood transcriptomes. (A) Principal component analysis (PCA) of whole blood transcriptomes from SLE group (n=11) and HC group (n=10). (B) Volcano plots of the gene expression comparison between SLE and HC. The horizontal axis represents the log2 (fold change) and the vertical axis represents the -log10 (P-value). The red plots represent the selected DEGs with fold change >=2 and p<0.01. (C) GO terms enriched in SLE patients’ samples. Genes with p-values < 0.05 were selected as input and enriched terms with padj < 0.05 were selected.

Next, to understand if there were differences in gene signatures between patients with or without CMD, we conducted a DE analysis comparing SLE-CMD to SLE-non-CMD. Due to the considerable variability among the samples and coupled with the fact that the dataset size is relatively small, a direct comparison between SLE groups revealed only 14 DE genes at padj < 0.1 (table 1). To comprehensively understand if there were differences within whole blood transcriptomes between SLE-CMD, SLE-non-CMD, and HC groups, we employed the HC as reference and conducted a comparative analysis of commonly up regulated and down regulated genes between SLE-CMD and SLE-non-CMD. This generated datasets that comprised genes commonly upregulated or downregulated between SLE-CMD and SLE-non-CMD versus healthy control and genes uniquely up or down regulated in both patient subgroups (as shown by the Venn diagrams in figure 2A, B). Our investigation unveiled 36 genes that were consistently upregulated and 20 genes that were downregulated across both SLE-CMD and SLE-non-CMD groups when compared to the HC group (overlapping area of figure 2A and B and table 2). Furthermore, we identified 176 genes displaying unique expression patterns between SLE-CMD and HC (left hand area of Venn diagrams in figure 2A and B), along with 144 unique DE genes between SLE-non-CMD and HC (right hand area of Venn diagrams in figure 2A and B). Analyzing the DE genes in common between SLE-CMD and SLE-non-CMD, Gene Ontology (GO) and pathway enrichment analyses indicated that these genes are clearly associated with antiviral immune responses and were primarily IFN stimulated genes (Figure 2C).

**Table 1.**
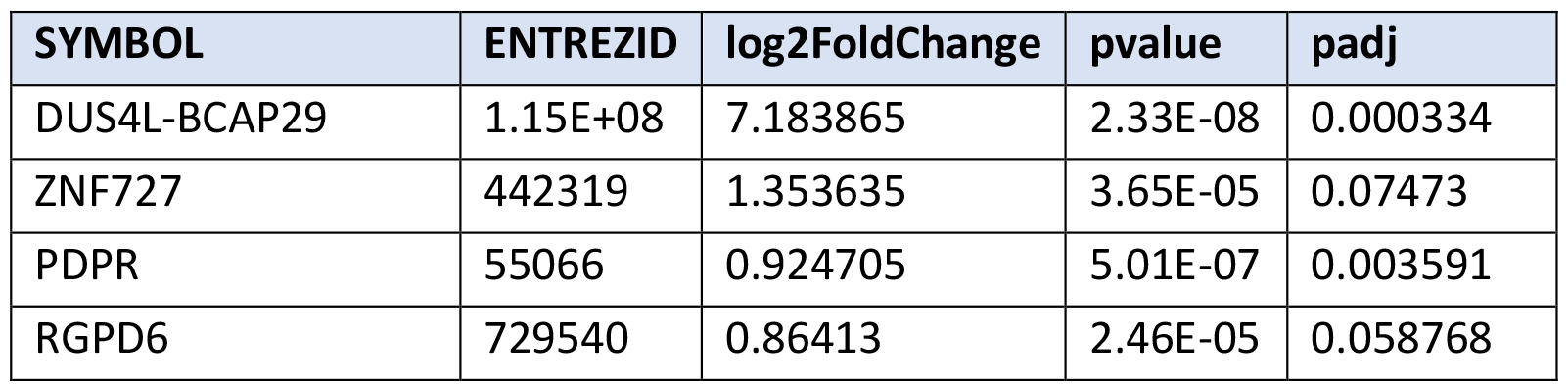
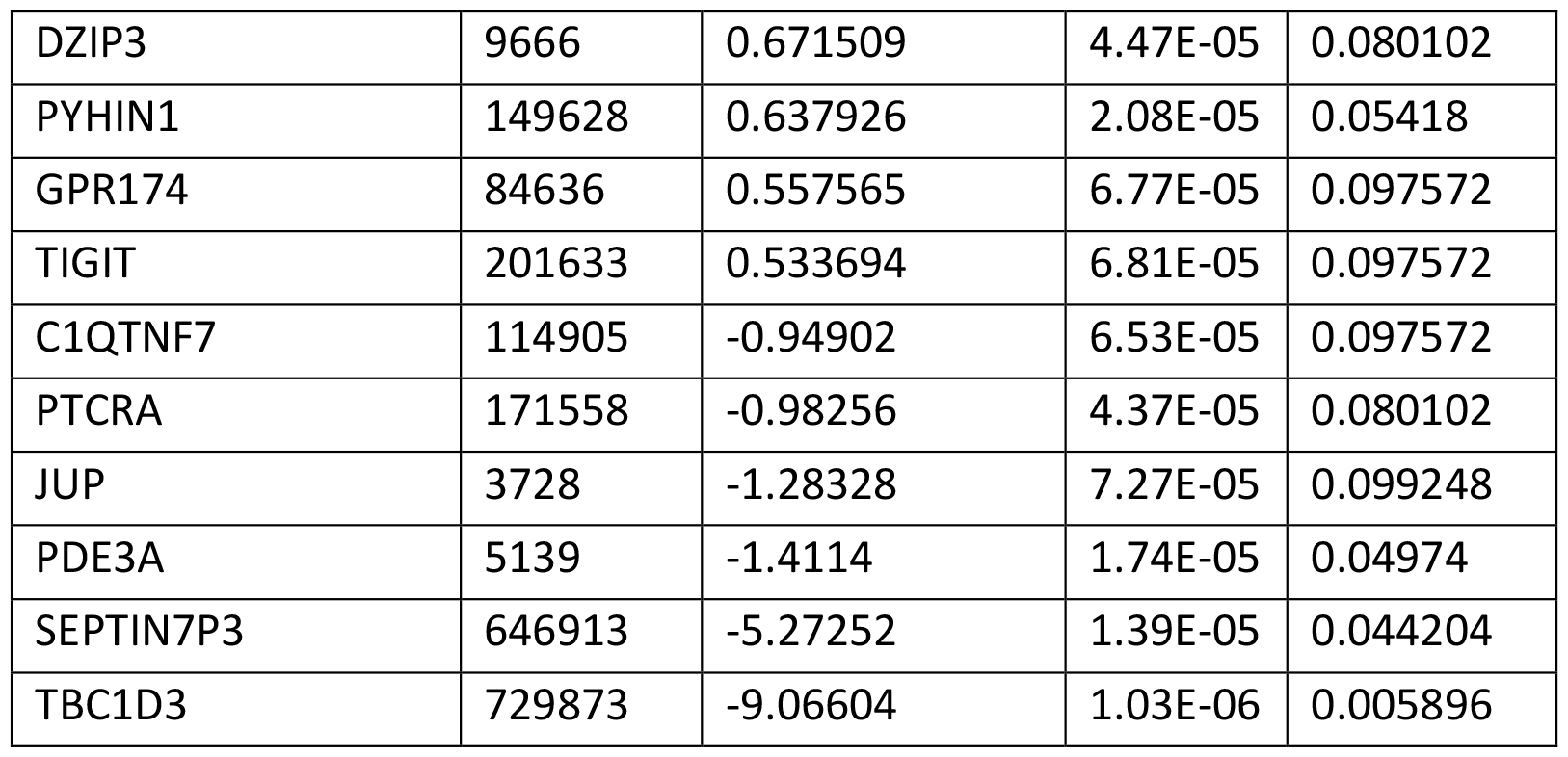
DEG between SLE-CMD and SLE-non-CMD. DE genes were identified using DEseq2 v1.42.0 with padj < 0.1. SLE-CMD (n=4) and SLE-non-CMD (n=7).

**Table 2.** Common genes between SLE-CMD vs HC and SLE-non-CMD vs HC. DE genes were identified using DEseq2 v1.42.0 with padj < 0.1, log2FoldChange > 0.5 as up regulated gene cutoff, and log2FoldChang < 0.5 as down regulated gene cutoff. SLE-CMD (n=4), SLE-non-CMD (n=7), and HC (n=10).

| Upregulated genes | Downregulated genes |
| --- | --- |
| SLC12A1 | TBX21 |
| SIGLEC1 | C1orf21 |
| OAS1 | RAB33A |
| OAS3 | COL6A2 |
| OAS2 | CEP78 |
| PRLR | GLB1L2 |
| IFIH1 | DLG5 |
| IFIT3 | PCDH1 |
| IFI6 | CLDND2 |
| XAF1 | SGCD |
| EPSTI1 | S1PR5 |
| RSAD2 | FCRL6 |
| CMPK2 | KLRC4 |
| RTP4 | TOGARAM2 |
| DDX60 | RN7SL653P |
| IFI44L | C19orf84 |
| IFI44 | H4C12 |
| HERC6 | RN7SL2 |
| HERC5 | RN7SL1 |

| NKD1 | RN7SL3 |
| --- | --- |
| ZCCHC2 |  |
| MX1 |  |
| LY6E |  |
| IFI27 |  |
| ANO5 |  |
| FBXO39 |  |
| USP18 |  |
| IFIT1 |  |
| ISG15 |  |
| PLSCR1 |  |
| SPATS2L |  |
| PGAP1 |  |
| GRASLND |  |
| CTXN2 |  |
| ZDHHC4P1 |  |
| CCDC194 |  |

**Figure 2.**
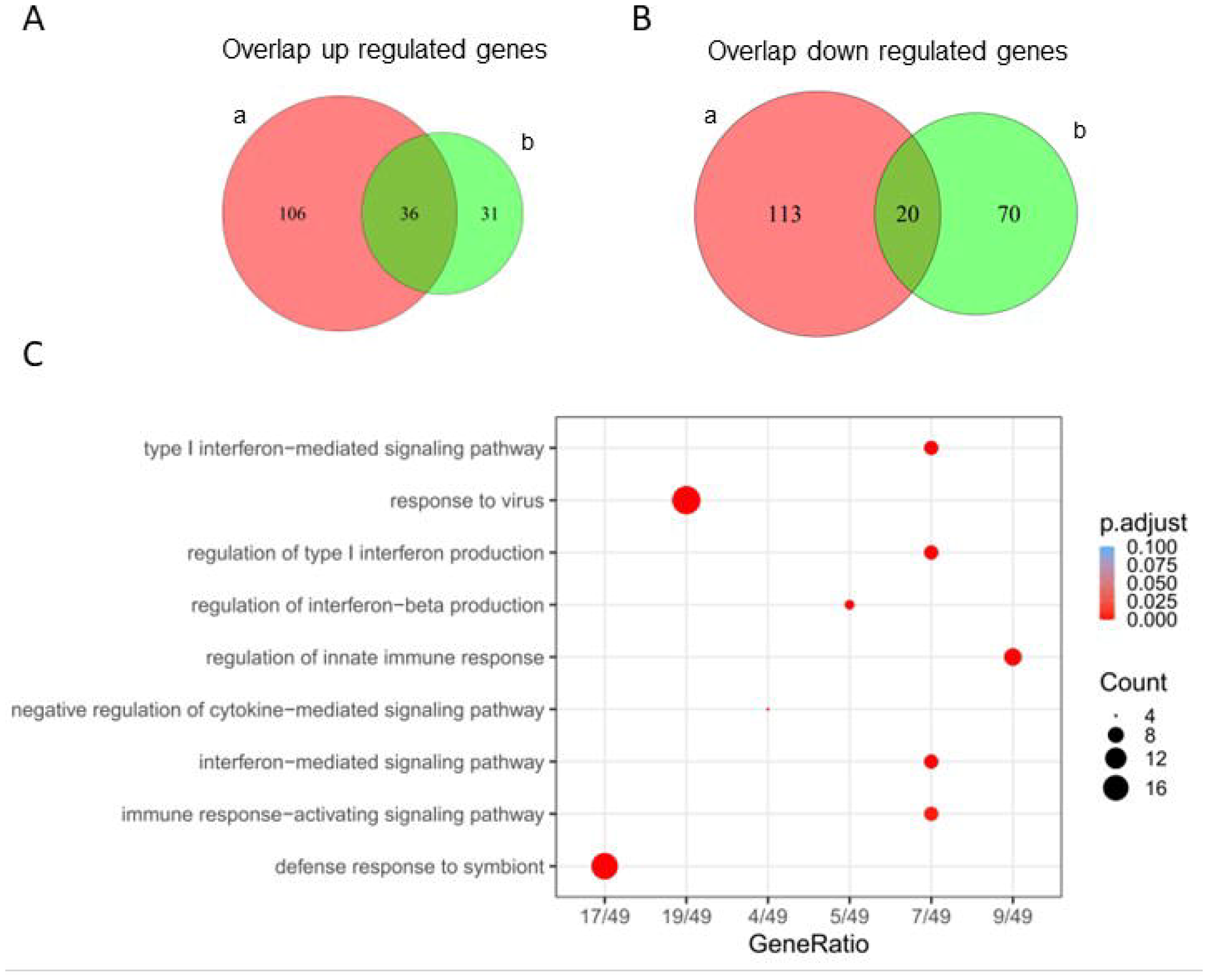
Unique gene differentially expressed in SLE-CMD and SLE-non-CMD whole blood samples using HC as reference group. (A, B) Venn diagram representing the unique or overlapping (A) upregulated or (B) downregulated genes in SLE-CMD and SLE-non-CMD when using HC as a reference group. SLE-CMD vs HC (a); SLE-non-CMD vs HC (b); DE genes were identified using DEseq2 v1.42.0 with padj < 0.1, log2FoldChange > 0.5 as up regulated gene cutoff, and log2FoldChang < 0.5 as down regulated gene cutoff. SLE-CMD (n=4), SLE-non-CMD (n=7), and HC (n=10). (C) Go pathway analysis for SLE-CMD and SLE-non-CMD common genes. 60 common genes were used for this test with default filters of pvalueCutoff = 0.05, qvalueCutoff = 0.2, minGSSize = 10, maxGSSize = 500.

We next conducted Ontology (GO) and pathway enrichment analyses for genes unique to SLE-CMD and SLE-non-CMD patients. Sorting by P value, the top GO terms of the biological process (BP), cellular component (CC) and molecular function (MF) categories are shown in Figure 3A (SLE-CMD) and 3B (SLE-non-CMD). As GO terms for RNA sensing, double stranded (ds) RNA and single stranded (ss) RNA binding, were enriched in SLE-CMD patient samples, we further analyzed the expression of the leading edge genes in the top GO terms (Figure 3C). In contrast only downregulated genes in SLE-non-CMD patients were associated with any GO categories. For example, genes associated with blood coagulation, cell-cell junction, and cellular defense response were decreased in SLE-non-CMD blood samples (Figure 3D).

**Figure 3.**
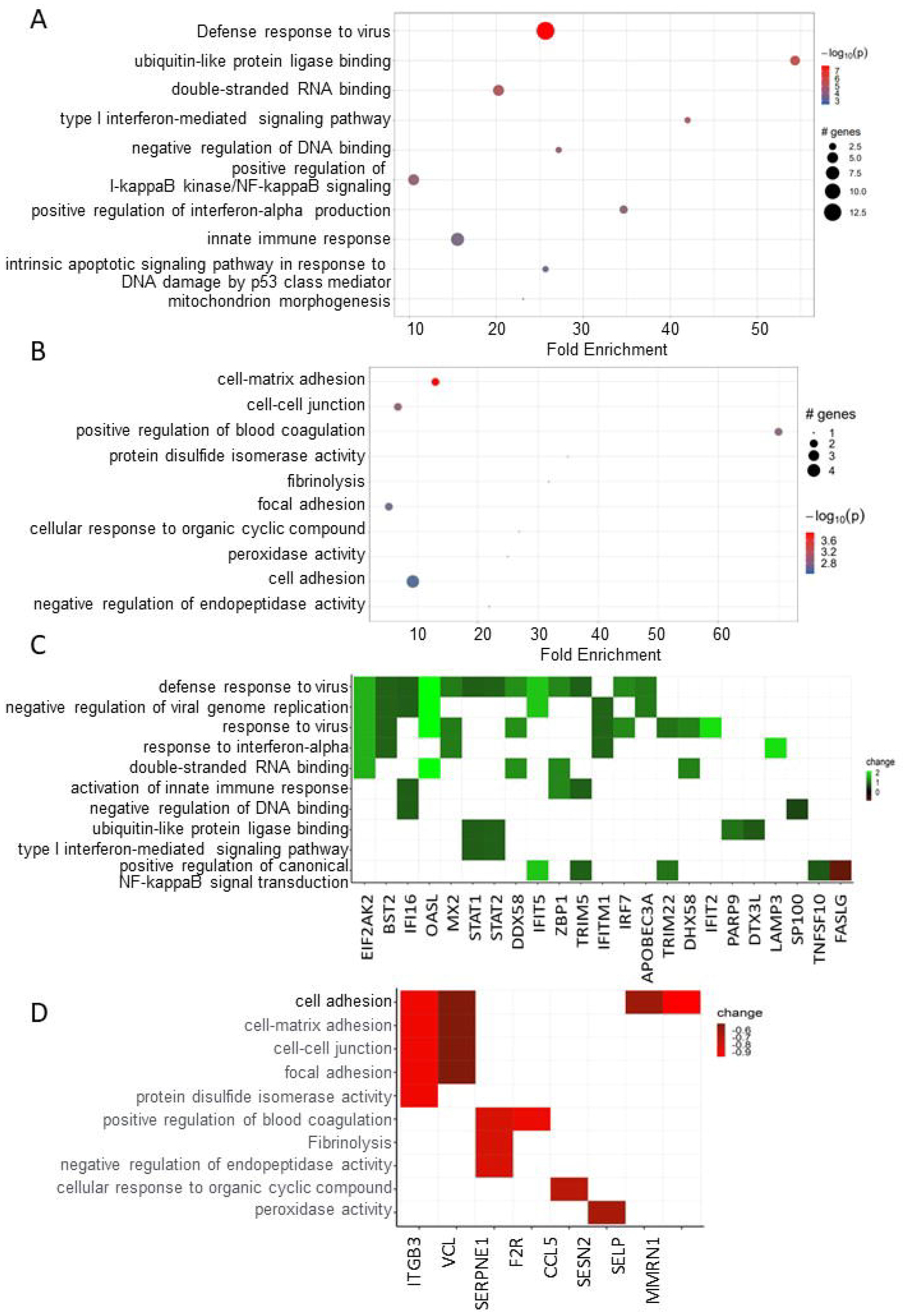
Pathway analysis of DE genes unique to CMD or non-CMD SLE patients. (A, B) Go pathway analysis for unique genes in (A) SLE-CMD and (B) SLE-non-CMD. Genes with p-values < 0.05 were selected as input and enriched terms with padj < 0.1 were selected. SLE-CMD (n=4), SLE-non-CMD (n=7), and HC (n=10). (C) Heatmap of SLE-CMD unique genes in the enriched GO terms. Green: up regulated; Red: down regulated. (D) Heatmap of unique genes in SLE-non-CMD in the enriched GO terms. Green: up regulated; Red: down regulated.

Analyzing a subset of genes relevant to RNA sensing, we observed that a panel of genes associated with IFN signaling and response to RNA and DNA sensing such as RIGI, DDX60, DHX58 and ZBP1 were significantly increased in SLE-CMD compared to HC and SLE-non-CMD (Figure 4A, B). In contrast, down regulated genes included the inhibitory receptor TIGIT, and markers of natural killer cells (NK) or invariant NK T cells (iNKT) KLRG1 and KLRC, suggesting that cells and pathways that might restrain cardiotoxic responses are decreased in CMD patient blood samples. Another interesting finding was the observation that IFNLR1, a component of the receptor for IFN lambda, was decreased in SLE-CMD compared to SLE-non-CMD and HC. Enzymes involved in ADP-ribosylation, a post-translational modification that regulates protein function, are also differentially upregulated in SLE-CMD patients compared to non-CMD. PARP9 and PARP14 were both increased and are part of a sub family of ADP-ribosylation enzymes that recognize mono-ADP-ribosylation (MAR) on proteins as opposed to poly-ADP-ribosylation (PAR). In patients with non-CMD, pathways relative to coagulation, platelet activity and cell adhesion are decreased specifically relative to SLE-CMD patients. These results suggest that RNA sensing pathways are associated with development of vascular changes associated with CMD in SLE, whereas reduced platelet activity and coagulation is associated with the reduced left ventricular function we observed in the non-CMD cohort in our previous cardiac MRI study [9].

**Figure 4.**
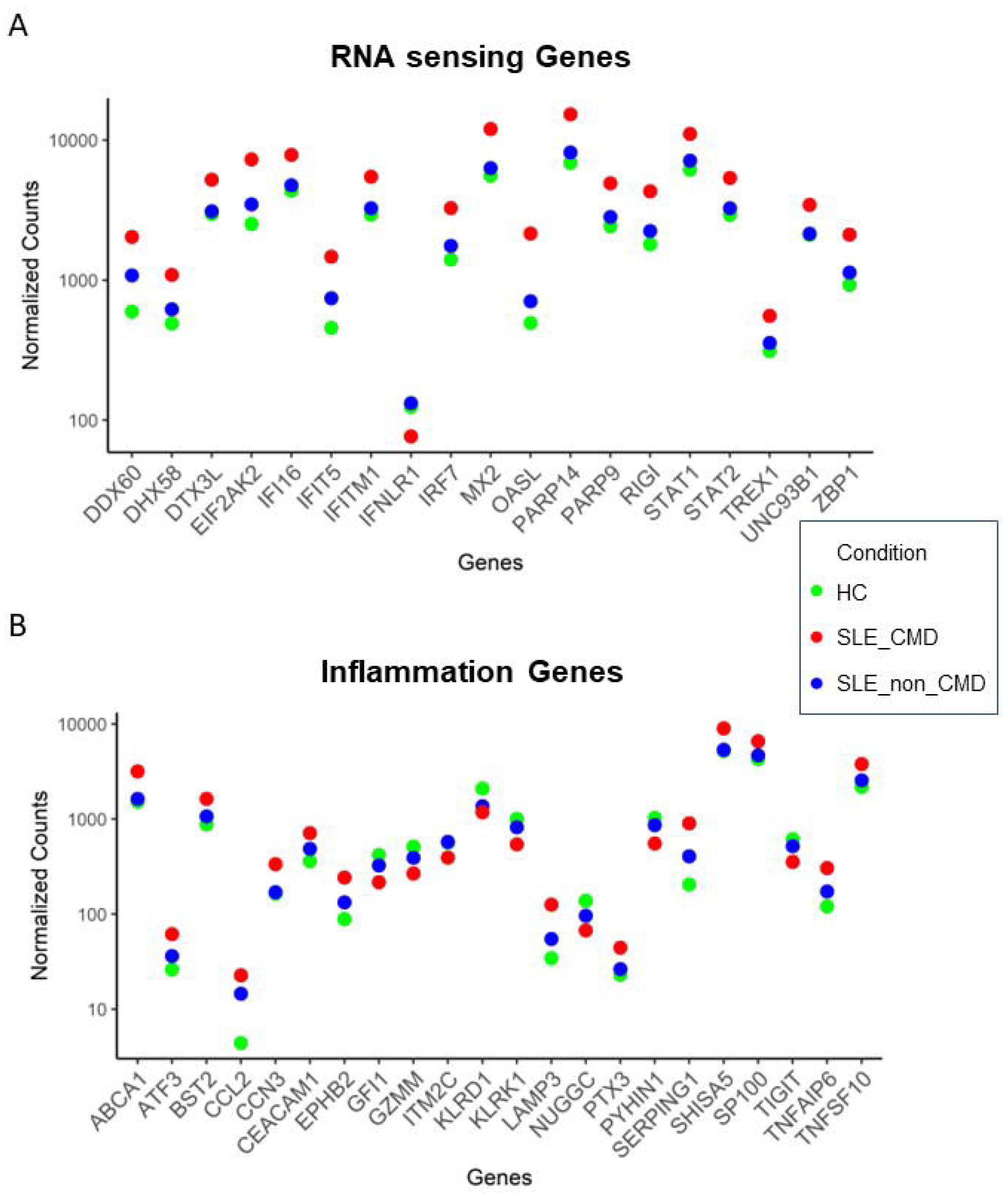
SLE-CMD unique gene signature related to RNA sensing and inflammation. (A,B) Dot plot of enrichment of (A) RNA sensing related genes and (B) inflammation related genes in SLE-CMD patient samples (Padj < 0.1, |log2FoldChange| > 0.5). SLE-CMD (n=4), SLE-non-CMD (n=7), and HC (n=10).

## Discussion

Chest pain is a common complaint among individuals with SLE even when there is no evidence of obstructive coronary artery disease (CAD)[8]. It is crucial to consider ischemic mechanisms like Coronary Microvascular Dysfunction (CMD) and coronary vasospasm during the diagnostic assessment. Given the heightened risk of cardiovascular disease (CVD) mortality and morbidity associated with SLE, stemming from inflammatory and metabolic pathophysiological processes, there is a pressing need for early identification and prevention of CVD risk factors. Hence, comprehending the molecular patterns within SLE-CMD may offer valuable insights into diagnostic methods and potential treatment approaches for clinical applications.

Endothelial dysfunction is a key driver of microvascular dysfunction and CMD. Endothelial cells (ECs) serve as a barrier between circulating blood components and the vascular wall and as such, play an important role in regulating immune responses to danger associated (such as oxLDL, heatshock proteins) and pathogen-associated molecular patterns (bacterial and viral proteins or nucleic acids) (DAMPs and PAMPs, respectively). In doing so they secrete cytokines and chemokines that regulate inflammation and immune responses, in addition to proteins that regulate vascular tone and function. Viral infection of ECs results in the activation of anti-viral signaling pathways that are activated to contain and eliminate the infection. One of the keyways viruses are detected by an infected cell is through recognition of viral RNA and DNA in the cytosol by RNA and DNA sensing proteins. RIG-I, MDA-5 and DHX58 (or LGP2) are three members of the RIG-I like receptors (RLRs) that are RNA helicases that recognize viral-derived dsRNA, become activated and drive the production of type I IFNs. A number of cardiotoxic agents such as 25-hydroxylcholesterol, angiotensin II and type I IFNs themselves have been shown to upregulate RIG-I expression. Together the increased expression of IFNs and IFN stimulated genes such as ISG15, CXCL10 and OAS2, are known to trigger EC dysfunction, specifically nitric oxide (NO) decreases, leukocyte adhesion, inflammation, coagulation, and endothelial cell injury. However, the direct role of RIG-I-mediated IFN induction in endothelial cells is relatively unexplored. One study showed that systemic administration of a RIG-I ligand led to impaired endothelium-dependent vasodilation in mouse aorta, although whether the effects of RIG-I activation were endothelial specific in this case was not shown. Our data clearly shows an association between upregulation of RIG-I, DHX58 and other components of RNA/DNA sensing such as ZBP2, in whole blood RNA samples from SLE patients with CMD compared to either healthy control subjects or SLE patients with chest pain but no evidence of CMD or obstructive disease. This data would suggest that increased levels of these enzymes systemically may result in increased secretion of ISGs such as OAS2 and CXCL10 that can alter endothelial function and potentially contribute to CMD. In addition, the increased expression of adhesion markers in these samples (CEACAM1 and X) would also suggest that the immune cells present are activated and will adhere to endothelial cells to a greater extent.

Another interesting and important finding in our study highlights the elevated levels of NKG2D, a marker associated with natural killer cells. NKG2D is a protein encoded by the KLRK1 gene, located within the NK-gene complex (NKC) on chromosome 6 in mice and chromosome 12 in humans[16]. It is expressed by various immune cells, including NK cells, γδ T cells, and CD8+ αβ T cells in humans[17, 18]. When NKG2D interacts with its ligands, it triggers or enhances the activity of immune cells expressing NKG2D. This activation prompts immune cells to proliferate, release cytokines (such as IFN-γ, granulocyte-macrophage colony-stimulating factor, macrophage inflammatory protein 1-α, and interleukin-2), and/or target and destroy cells expressing the ligands[19]. The heightened presence of NKG2D ligands in aortic plaques suggests a potential role in immune activation associated with atherosclerosis[20]. Notably, studies have indicated that the interaction between immune cells and cardiomyocytes via NKG2D/NKG2DL can induce cardiomyocyte death, exacerbating cardiac remodeling following myocardial infarction (MI)[21]. This underscores the importance of understanding NKG2D’s involvement in cardiovascular pathologies and its potential as a target for therapeutic interventions.

The reduction of the T cell specific inhibitory receptor TIGIT distinguished SLE-CMD from SLE-non-CMD whole blood samples. TIGIT has previously been shown to be elevated on SLE T cells which has prompted the development of TIGIT-targeting strategies for SLE. Here our results indicate that in contrast TIGIT levels are decreased and correlate with SLE-CMD. Exhausted T cells express TIGIT and are not only a feature of tumor immunity but also of SLE and lupus nephritis in the MRL/lpr model of SLE[22]. However, the relevance of this to human disease has been questioned from scRNAseq analyses by the AMP consortium[23]. Our finding in SLE-CMD that TIGIT levels are decreased suggests that there is less of an exhausted phenotype in our cohort of SLE-CMD compared to SLE-non-CMD patients and that potentially T cell activity is contributing to microvascular defects.

PARP9 and PARP14 have previously been shown to positively regulate signaling via RNA sensing pathways and IFNβ induction. In the case of PARP14, it is thought to play a role in epigenetic regulation of gene expression. PARP9 on the other hand has been described as a non-canonical RNA sensor that binds viral RNA and promotes PI3K/AKT activation, leading to IRF3 and IRF7 phosphorylation and IFNβ induction in mouse dendritic cells[24-26]. Interestingly PARP14 also has an RNA binding domain, although whether this plays a role in IFN responses is unknown.

In conclusion this study has unveiled a RNA sensing gene signature in whole blood samples of SLE patients with CMD compared to SLE patients without. Whilst our study is limited by the small samples size our signature is robust and agrees with recent literature examining the role of RIG-I and TREX1 in heart disease. While circulating microRNAs have been suggested as biomarkers for early-stage CAD, to date no biomarkers for CMD have been described. Both oxidative stress and epigenetic regulation of such pathways have previously been linked with CMD. Whether mitochondrial function and oxidative stress are contributing to mtRNA and DNA release in SLE with CMD patient immune cells is a potential mechanism driving these findings [27, 28]. Interestingly, CMD is common in patients with prior COVID infection and CMD considered a strong driver of morbidity and mortality associated with COVID-19 **[29, 30]**. Other viral infections such as HIV have also been linked to CMD in infected individuals without risk factors for CVD [31]. Thus, this opens up many questions with respect to enhanced risk of COVID-19-driven complications in patients with autoimmune disease and specifically SLE. Further analysis is required to understand the relationship between this gene signature, CMD and immune cell dysregulation.

